# Changes in serial sarcomere number of five hindlimb muscles across adult aging in rats

**DOI:** 10.1101/2025.03.04.641485

**Authors:** Avery Hinks, Geoffrey A. Power

## Abstract

**Introduction:** Aging is associated with a reduction in muscle fascicle length (FL), which contributes to sarcopenia: the age-related loss of muscle mass and function. Studies on rodents have confirmed this reduced FL is driven by a loss of sarcomeres aligned in series (serial sarcomere number; SSN) along a muscle. However, studies on rodents have focused primarily on rat plantar flexor SSN at two aging timepoints, leaving an incomplete view of age-related changes in SSN. Hence, this study investigated SSN as a contributor to the age-related loss of muscle mass in five hindlimb muscles across four aging timepoints in rats.

**Methods:** The soleus, medial gastrocnemius (MG), plantaris, tibialis anterior (TA), and vastus lateralis (VL) were obtained from 5 young (8 months), 5 middle-aged (20 months), 5 old (32 months), and 5 very old (36 months) male F344BN rats. After fixation of muscles in formalin and digestion in nitric acid, fascicles were teased out end-to-end to measure FL. SSN was determined by dividing FL by sarcomere length measured via laser diffraction. Muscle wet weight, anatomical cross-sectional area (ACSA), and physiological cross-sectional area (PCSA) were also determined for insight on age-related losses of whole-muscle mass and in-parallel muscle morphology.

**Results:** Age-related SSN loss was apparent after middle age for all muscles, with the plantaris showing the smallest (8%) and the VL the greatest (21%) loss. The MG and VL appeared to plateau in their SSN loss by 32 months, while the soleus and TA underwent continued loss from 32 to 36 months. In all muscles, SSN loss evidently contributed in part to the loss of muscle mass, alongside losses of contractile tissue in parallel (indicated by ACSA and PCSA).

**Conclusion:** As SSN is closely tied to biomechanical function, these findings present SSN as a distinct target for improving muscle performance in older adults.

## Introduction

Aging is associated with a reduction of muscle fascicle length (FL) (1,2). Studies on rodents have shown this reduction of FL is driven by a loss of sarcomeres aligned in series (serial sarcomere number; SSN) (3–6). This age-related loss of SSN has deleterious effects on muscle mechanical performance including a leftward shift in the torque-angle relationship, reduced power production, and elevated passive stiffness (5,6). Current techniques for measuring SSN in human muscle are invasive or limited (7,8), therefore animal models—in which muscles are dissected off the limb for measurements of sarcomere length (SL) via laser diffraction—are necessary for understanding how SSN changes with age. Currently, studies on rodents have focused primarily on rat plantar flexor muscles (4–6), limiting applicability to other muscle groups. Furthermore, these studies have only investigated two aging time points (i.e., young 8-month-old rats vs. old 32-month-old rats), leaving an incomplete view of changes in SSN throughout adult aging.

The loss of muscle mass with age could be attributed to losses of both sarcomeres aligned in series and in parallel across a muscle (9,10). Additionally, in muscles with a greater fascicle pennation angle, age-related decreases in anatomical cross-sectional area (ACSA) (a well-known hallmark of aging muscle (11)) could also be driven by losses of serially aligned sarcomeres rather than just in parallel (9). However, it is currently unclear to what extent the age-related loss of SSN contributes to the loss of overall muscle mass and consequently function. Investigating age-related losses of SSN and physiological cross-sectional area (PCSA; an indicator of contractile tissue in parallel at the whole-muscle level) across multiple muscles would further current understanding of where muscle contractile tissue is lost (i.e., in parallel vs. in series) with age and implications on muscle contractile performance.

The purpose of this study was to investigate age-related SSN loss in F344BN rats among plantar flexor, dorsiflexor, and knee extensor muscles across adulthood. Timepoints included: young adult (8 months; ∼25 “human years”), middle-aged (20 months; ∼50 “human years”), old (32 months; ∼80 “human years”), and very old (36 months; ∼90 “human years”). Due to their role in ambulation and ease/consistency of dissection for obtaining intact muscle fascicles, we examined the soleus, medial gastrocnemius (MG), plantaris, tibialis anterior (TA), and vastus lateralis (VL).

## Methods

### Animals

Five young (5 months), 5 middle-aged (20 months), 5 old (32 months), and 5 very old (36 months) male Fischer 344/Brown Norway F1 rats were obtained (Charles River Laboratories, Senneville, QC, Canada). These rat ages correspond to human ages of approximately 25, 50, 80, and 90 years, respectively (12). Approval was obtained from the University of Guelph’s Animal Care Committee (AUP #4905) and all protocols followed CCAC guidelines. Rats were housed at 23°C in groups of two or three and given ad-libitum access to a Teklad global 18% protein rodent diet (Envigo, Huntington, Cambs., UK) and room-temperature water. Rats were allowed at least 1 week to acclimate to their housing conditions, then they were euthanized via isoflurane followed by CO_2_ asphyxiation and cervical dislocation.

### Muscle fixation

The left hindlimb was amputated and fixed in 10% phosphate-buffered formalin with the ankle, knee, and hip pinned at 90°. After fixation for 1-2 weeks, the soleus, MG, plantaris, TA, and VL (Figure 1A) were dissected and rinsed with phosphate-buffered saline then transferred back to formalin for at least 1 additional week of fixation. Muscles were weighed following this final fixation step.

**Figure 1:**
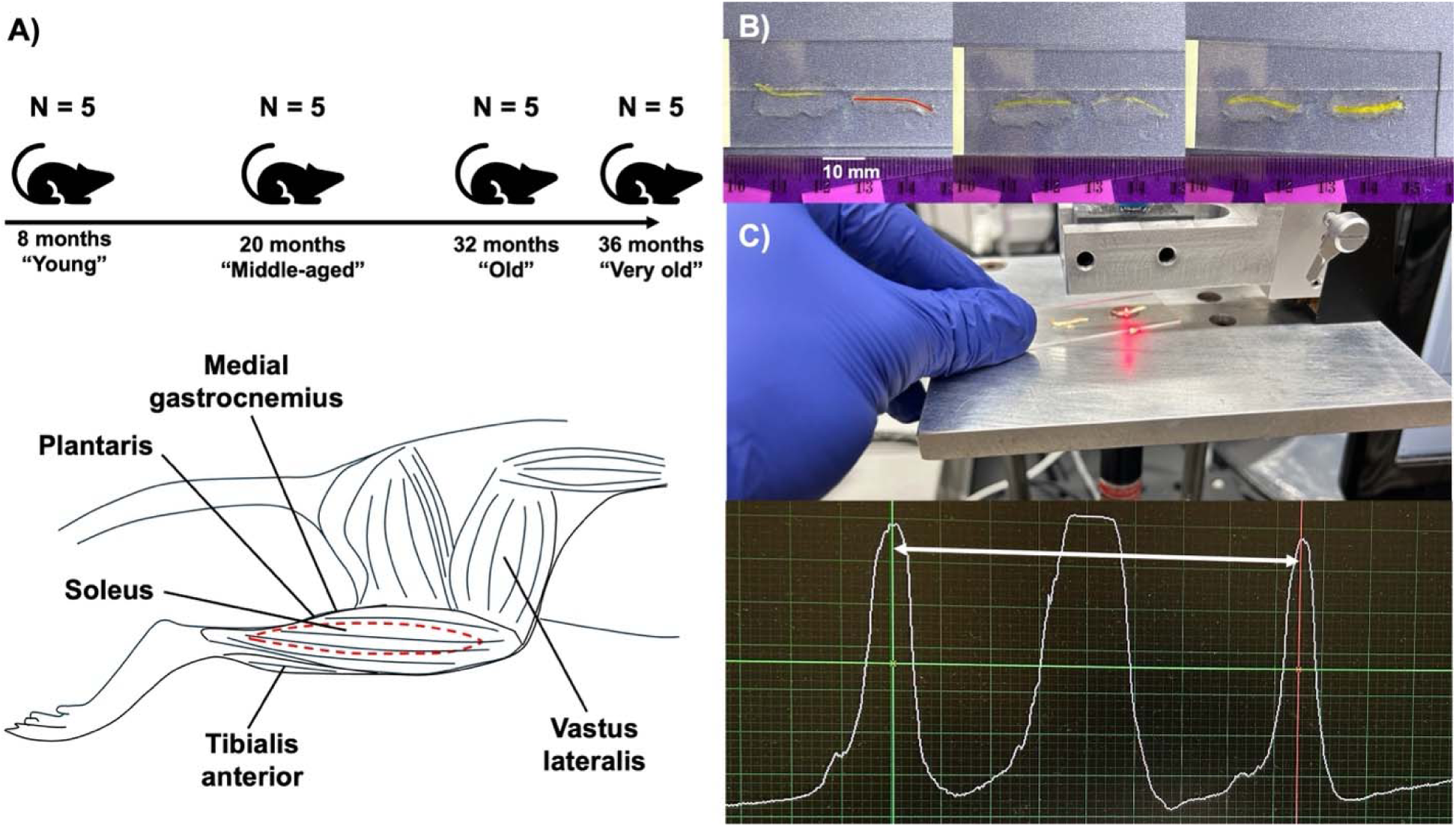
**A.** Five young (8 months), 5 middle-aged (20 months), 5 old (32 months), and 5 very old (36 months) rats were obtained from the National Institutes on Aging. The soleus (deep, dashed red lines), medial gastrocnemius (medial lower hindlimb, not pictured), plantaris (posterior lower hindlimb, not pictured), tibialis anterior, and vastus lateralis were obtained. **B.** Fascicle lengths were measured end-to-end (demonstrated by the red line) in ImageJ. **C.** Sarcomere lengths were measured in each fascicle using laser diffraction, with the distance between the 1^st^ order peaks used to calculate sarcomere lengths.

### Muscle anatomical cross-sectional area

Following fixation in formalin, the width (medial-lateral dimension) and thickness (anterior-posterior dimension) of the muscles were measured at the mid-muscle belly using calipers. ACSA was then determined using the equation:

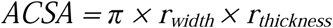

With *r_width_* and *r_thickness_* being the radii in the width and thickness dimensions, respectively (i.e., width or thickness divided by 2). This equation assumes ACSA is elliptical (13).

### Serial sarcomere number determinations

The muscles were removed from formalin, rinsed with phosphate-buffered saline, then digested in 30% nitric acid for 6-8 hours to remove connective tissue and allow for individual muscle fascicles to be teased out (14,15).

For each muscle, two fascicles were obtained from each of the proximal, middle, and distal regions of the muscle (i.e., n=6 fascicles total per muscle). Dissected fascicles were placed on glass microslides (VWR International, USA), then FLs were measured using ImageJ software (version 1.53f, National Institutes of Health, USA) from pictures captured by a level, tripod-mounted digital camera, with measurements calibrated to a ruler in plane with the fascicles (Figure 1B). Sarcomere length measurements were taken at six different locations proximal to distal along each fascicle via laser diffraction with a 5-mW diode laser (∼1 mm beam diameter, 635 nm wavelength; Coherent, Santa Clara, CA, USA) and custom LabVIEW program (Version 2011, National Instruments, Austin, TX, USA) (16) (Figure 1C). Given the laser diameter of ∼1 mm, one SL measurement itself represents an average of hundreds to thousands of SLs. Our total quantity of SL and FL measurements is consistent with previous studies (14,15,17). Within each fascicle, the six SL measurements were averaged to obtain a value of average SL. Serial sarcomere number of each fascicle was calculated as:

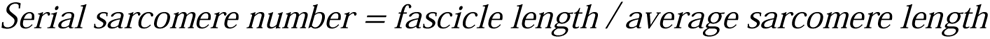

### Determination of physiological cross-sectional area

To gain further insight on changes in contractile tissue in parallel at the whole muscle level, we calculated physiological cross-sectional area (PCSA) using the equation (13,17,18):

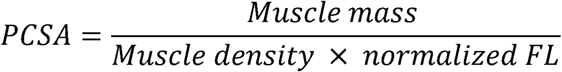

Muscle density was assumed to be 1.112 g/cm^3^ (18). Normalized FL was calculated using FL of dissected fascicles and measured SL in the equation (18):

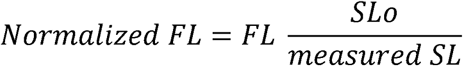

SLo represents resting SL of rat skeletal muscle at the optimal muscle length, assumed to be ∼2.7 μm based on previous literature (17,19).

### Statistical analyses

In SPSS Statistics Premium 28, normality of data was confirmed using Shapiro-Wilk tests. One-way analysis of variance (ANOVA) was used to investigate differences in body mass between age groups. Two-way ANOVA (age × muscle) was used to investigate muscle-dependent differences across age groups for muscle wet weight, ACSA, and PCSA. To analyse muscle-dependent age-related differences in FL, average SL, and SSN, we used a three-way nested ANOVA, including subject as a factor and viewing each fascicle separately but nested within each muscle (20). A Greenhouse-Geisser correction was applied for all ANOVAs if the sphericity assumption was not met. A Sidak correction was applied to post-hoc pairwise comparisons. Significance was set at α=0.05.

## Results

### Exclusion

One plantaris muscle from a middle-aged rat was damaged during the dissection process and was thus excluded from the study (i.e., data from n=4 middle-aged plantares are reported).

### Body mass

There was an effect of age on body mass (*P*=0.005) (Figure 2). Body mass was greater in middle-aged (+27%, *P*=0.004) and old (+21%, *P*=0.034) compared to young rats, but tapered off thereafter such that body mass did not differ between young and very old rats (*P*=0.275).

**Figure 2:**
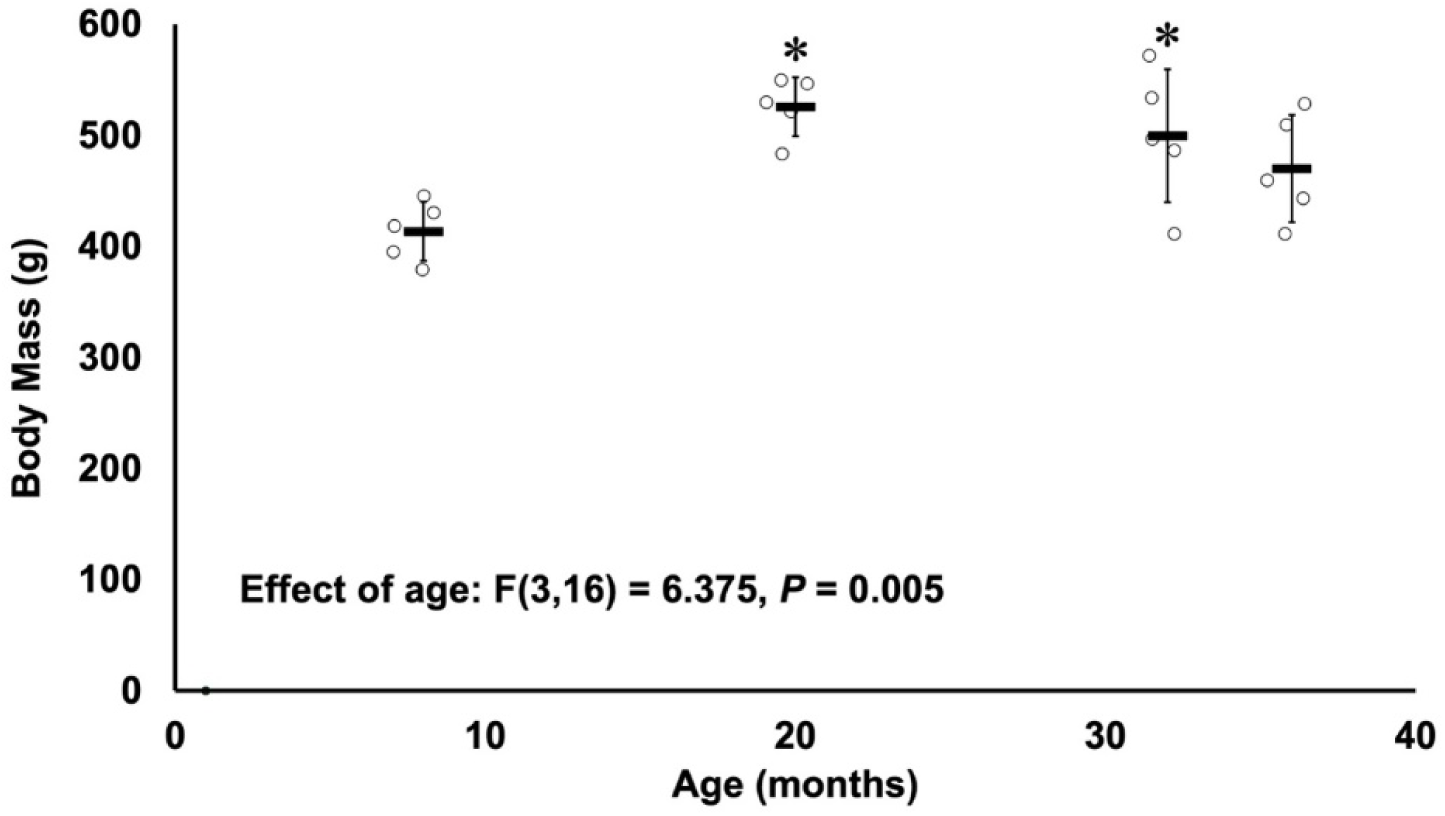
Changes in body mass across young (8 months), middle-aged (20 months), old (32 months), and very old (36 months) rats (n=5 per group). *Difference from young (P<0.05).

### Muscle-dependent changes in morphology across age groups

There were age × muscle interactions for muscle wet weight, ACSA, PCSA (Figure 3), FL, SL, and SSN (Figures 3-5) (all *P*<0.001), indicating that age-related changes in these variables were muscle dependent. Hence, age-related changes for each muscle are separated into their own sections below for post-hoc comparisons across age groups.

**Figure 3:**
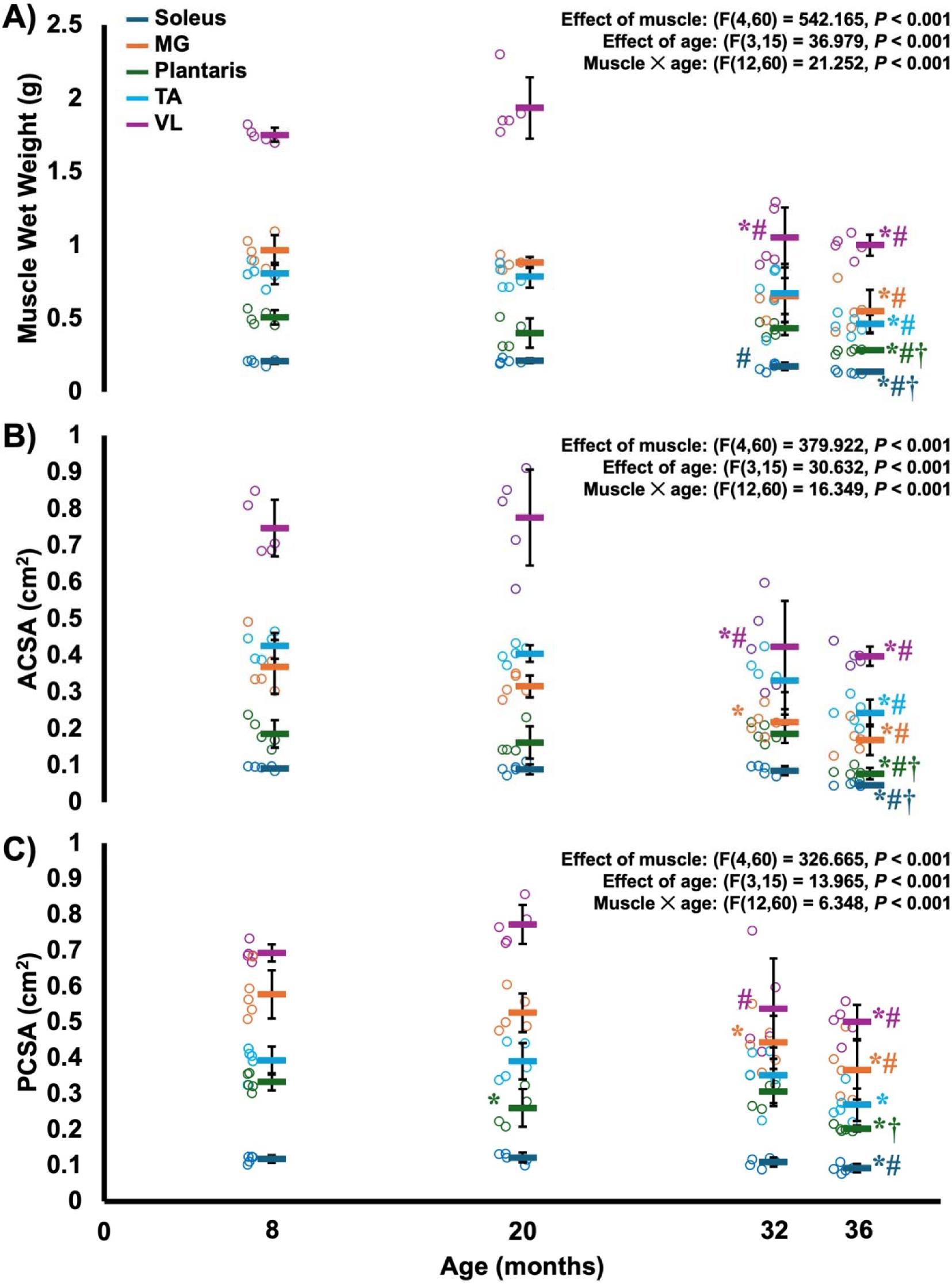
Changes in muscle wet weight **(A)**, anatomical cross-sectional area (ACSA) **(B)**, and physiological cross-sectional area (PCSA) **(C)** across young (8 months), middle-aged (20 months), old (32 months), and very old (36 months) rats in the soleus, medial gastrocnemius (MG), plantaris, tibialis anterior (TA), and vastus lateralis (VL). *Different from young (*P*<0.05). #Different from middle-aged (*P*<0.05). †Different from old (*P*<0.05). n=5 per group, except for the plantaris of middle-aged rats, which was n=4.

### Changes in soleus morphology across age groups

Soleus wet weight did not differ between young and middle-aged rats (*P*=0.997), but decreased 18% from middle-aged to old (*P*=0.033) and 21% from old to very old rats (*P*=0.038) with an overall decrease of 35% (Figure 3A).

Soleus ACSA did not differ among young, middle-aged, and old rats (*P*=0.918-0.999), but decreased in very old rats compared to all other age groups (all *P*<0.001) with the largest decrease of 50% from young to very old (Figure 3B). Similarly, soleus PCSA did not differ among young, middle-aged, and old rats (*P*=0.657-0.999), but decreased from young and middle-aged to very old rats (*P*=0.019-0.029) with the largest decrease of 24% from middle-aged to very old (Figure 3C).

Soleus FL as measured at a neutral joint angle did not differ between young and middle-aged rats (*P*=0.454), but decreased 11% from young to old (*P*<0.001) and 16% from young to very old (*P*<0.001), with a 6% decrease from old to very old (*P*=0.049) (Figure 4A). Soleus SL as measured at a neutral joint angle displayed some changes across ages, decreasing 4% from young to middle-aged rats (*P*=0.001) then increasing again from middle-aged to old rats (*P*=0.017) such that old (*P*=0.965) and very old (*P*=0.940) SL did not differ from young SL (Figure 5A). With SL staying constant between young, old, and very old rats, the decreases in FL were driven by losses of SSN. Specifically, soleus SSN decreased 10% from young to old rats (*P*<0.001) and 15% from young to very old rats (*P*<0.001) (Figure 6A). Thus, there was a progressive loss of soleus SSN after middle age.

**Figure 4:**
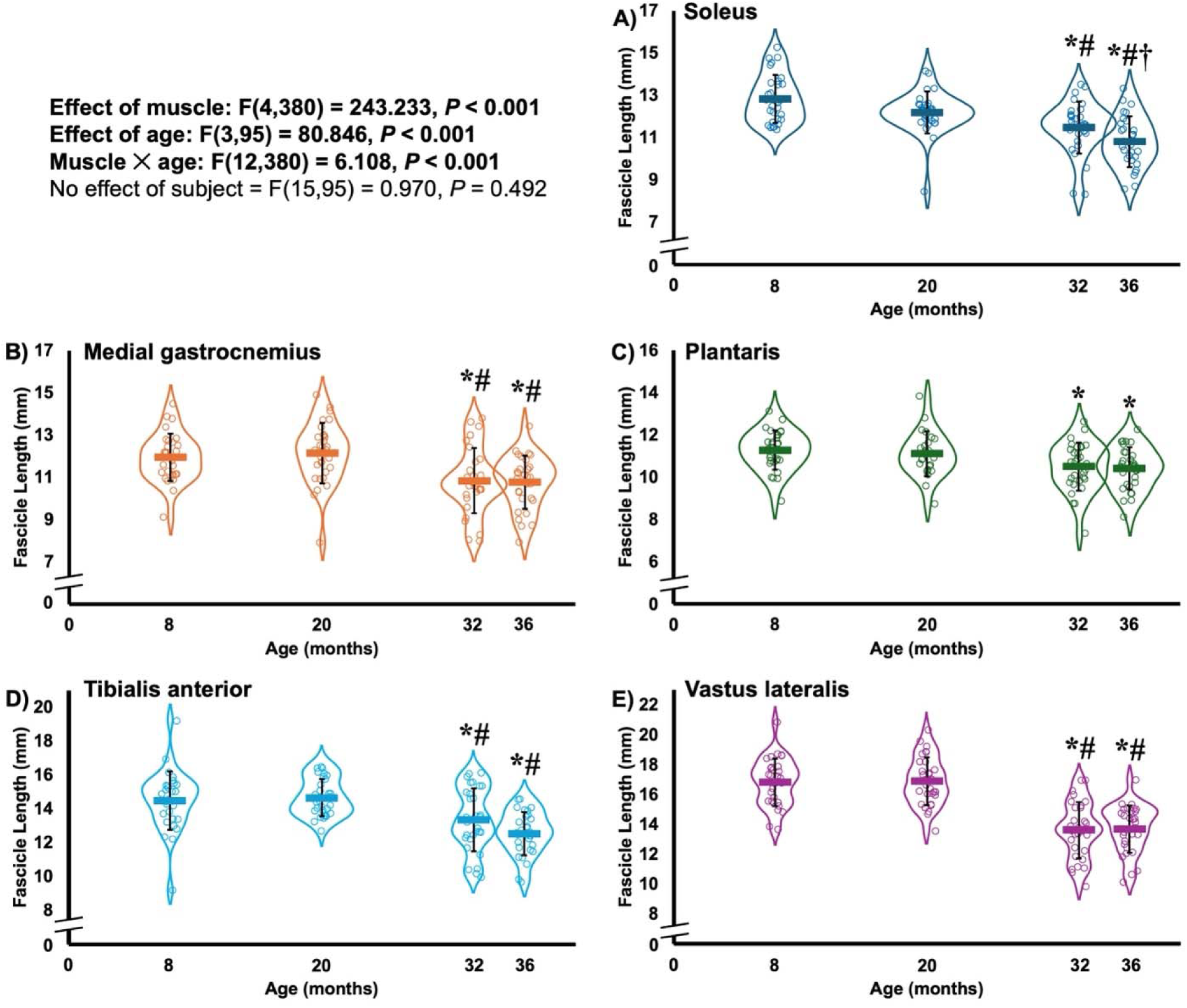
Violin plots showing changes in fascicle length across young (8 months), middle-aged (20 months), old (32 months), and very old (36 months) rats in the soleus **(A)**, medial gastrocnemius **(B)**, plantaris **(C)**, tibialis anterior **(D)**, and vastus lateralis **(E)**. *Different from young (*P*<0.05). #Different from middle-aged (*P*<0.05). †Different from old (*P*<0.05). Data from n=5 rats per group and n=6 fascicles per muscle (n=30 per group total) are displayed, except for the plantaris of middle-aged rats, which was n=4. Statistical analyses were run with fascicles nested within each rat (i.e., with an effect of subject included).

**Figure 5:**
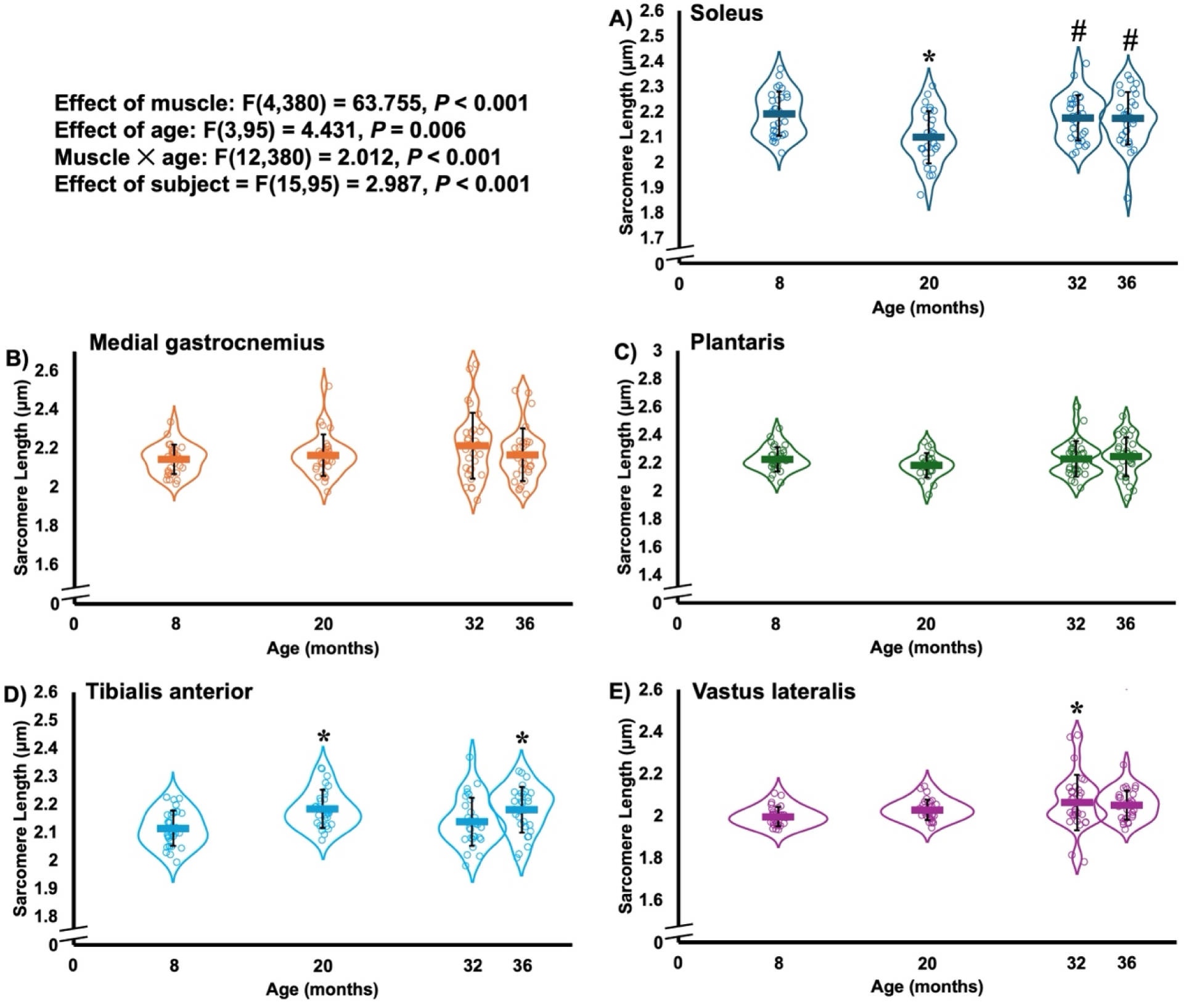
Violin plots showing changes in sarcomere length (i.e., mean sarcomere length within a fascicle) across young (8 months), middle-aged (20 months), old (32 months), and very old (36 months) rats in the soleus **(A)**, medial gastrocnemius **(B)**, plantaris **(C)**, tibialis anterior **(D)**, and vastus lateralis **(E)**. *Different from young (P<0.05). #Different from middle-aged (P<0.05). Data from n=5 rats per group and n=6 fascicles per muscle (n=30 per group total) are displayed, except for the plantaris of middle-aged rats, which was n=4. Statistical analyses were run with fascicles nested within each rat (i.e., with an effect of subject included).

**Figure 6:**
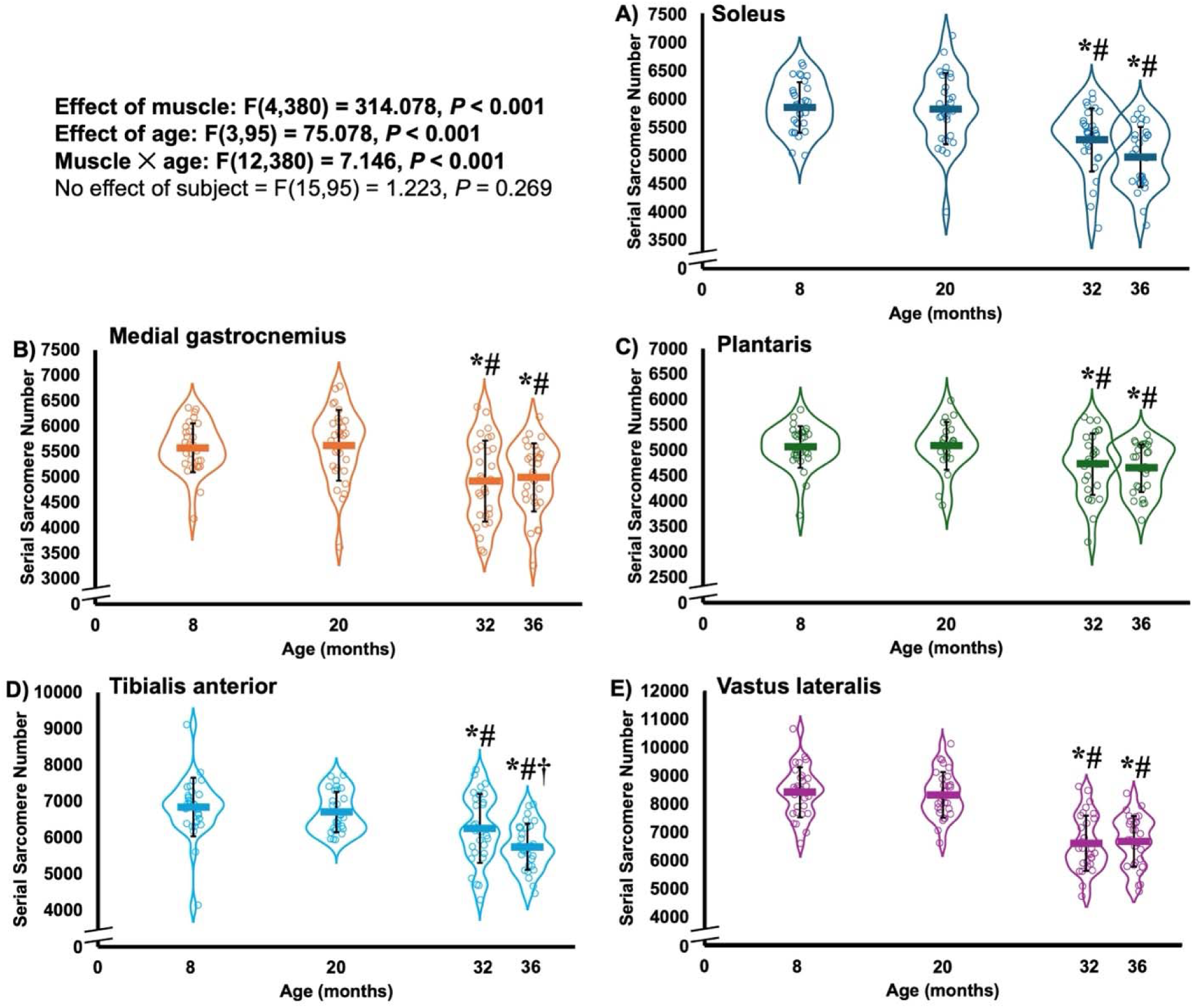
Violin plots showing changes in serial sarcomere number across young (8 months), middle-aged (20 months), old (32 months), and very old (36 months) rats in the soleus **(A)**, medial gastrocnemius **(B)**, plantaris **(C)**, tibialis anterior **(D)**, and vastus lateralis **(E)**. *Different from young (*P*<0.05). #Different from middle-aged (*P*<0.05). †Different from old (*P*<0.05). Data from n=5 rats per group and n=6 fascicles per muscle (n=30 per group total) are displayed, except for the plantaris of middle-aged rats, which was n=4. Statistical analyses were run with fascicles nested within each rat (i.e., with an effect of subject included).

Collectively, it appears that age-related losses of soleus muscle mass were driven mainly by losses of SSN at old age (32 months) and by both SSN and CSA at very old age (36 months).

### Changes in medial gastrocnemius morphology across age groups

MG wet weight did not differ between young and middle-aged rats (*P*=0.756) nor between old and very old rats (*P*=0.660), but decreased 32% from young to old (*P*=0.003) and 43% from young to very old (*P*<0.001) (Figure 3A).

MG ACSA did not differ between young and middle-aged rats (*P*=0.402) nor between old and very old rats (*P*=0.603), but decreased 41% from young to old (*P*=0.001) and 54% from young to very old (*P*<0.001) (Figure 3B). MG PCSA also did not differ between young and middle-aged rats (*P*=0.608) nor between old and very old rats (*P*=0.464), but decreased 23% from young to old (*P*=0.044) and 37% from young to very old (*P*=0.001) (Figure 3C).

MG FL as measured at a neutral joint angle also did not differ between young and middle-aged rats (*P*=0.739) nor between old and very old rats (*P*=1.00), but decreased from both young and middle-aged rats to old and very old rats (*P*<0.001-0.005) with an overall FL decrease of 11% (Figure 4B). MG SL as measured at a neutral joint angle did not differ across age groups (*P*=0.207-1.00) (Figure 5B), therefore the age-related decrease in MG FL was due to the loss of SSN. Similar to FL, MG SSN did not differ between young and middle-aged rats (*P*=0.937) nor between old and very old rats (*P*=0.999), but decreased from both young and middle-aged rats to old and very old rats (*P*<0.001-0.004) with an overall SSN loss of 12% (Figure 6B). Therefore, the MG lost SSN with age, but unlike the soleus, plateaued by 32 months.

Altogether, losses of MG muscle mass with age seemed to be driven by losses of both CSA and SSN.

### Changes in plantaris morphology across age groups

Plantaris wet weight did not differ among young, middle-aged, and old rats (*P*=0.077-0.964) but decreased 44% from young to very old (*P*<0.001), with a 34% decrease from old to very old (*P*=0.005) (Figure 3A).

Plantaris ACSA did not differ among young, middle-aged, and old rats (*P*=0.858-1.000), but decreased 58% from young to very old (*P*<0.001), and notably also decreased 58% from old to very old (*P*<0.001) (Figure 3B). Plantaris PCSA was 22% lower in middle-aged compared to young rats (*P*=0.035), though was not different between young and old rats (*P*=0.780), but still decreased 39% from young to very old (*P*<0.001) with also a 34% decrease from old to very old (*P*=0.001) (Figure 3C).

The plantaris saw less drastic age-related changes in FL and SSN than the soleus and MG. Plantaris FL did not differ between young and middle-aged rats (*P*=0.991) and only decreased 7% and 8% from young to old (*P*=0.021) and very old rats (*P*=0.008), respectively, with no difference between old and very old (*P*=1.00) (Figure 4C). Plantaris SL did not differ between any age groups (*P*=0.119-1.00) (Figure 5C), therefore the slight decrease in FL with age was attributed to a loss of SSN. Plantaris SSN did not differ between young and middle-aged rats (*P*=0.991) nor between old and very old rats (*P*=0.989), but decreased from both young and middle-aged rats to old and very old rats (*P*=0.007-0.045) with an overall SSN loss of 9% (Figure 6C). Therefore, like the MG, the plantaris lost SSN with age, plateauing by 32 months.

Altogether, the age-related loss of plantaris muscle mass, particularly observed in very old rats, looked to be primarily driven by losses of CSA and less so by losses of SSN.

### Changes in tibialis anterior morphology across age groups

TA wet weight did not differ among young, middle-aged and old rats (*P*=0.443-0.999) but decreased 43% from young to very old (*P*=0.002), although with no difference between old and very old (*P*=0.080) (Figure 3A).

TA ACSA also did not differ among young, middle-aged and old rats (*P*=0.098-0.976) but decreased 43% from young to very old (*P*<0.001), although with no difference between old and very old (*P*=0.128) (Figure 3B). Likewise, TA PCSA did not differ between young, middle-aged, and old rats (*P*=0.829-0.999) but decreased 31% from young to very old (*P*<0.001), although with no difference between old and very old (*P*=0.169) (Figure 3C).

The TA saw similar age-related changes as the soleus. TA FL did not differ between young and middle-aged rats (*P*=0.961), but decreased from both young and middle-aged rats to old and very old rats (*P*<0.001-0.018) with a 9% and 14% decrease from middle-aged to old and very old, respectively (Figure 4D). TA SL also exhibited some slight changes across age groups, increasing 3% from young to middle-aged rats (*P*=0.023), decreasing from middle-aged to old rats such that it did not differ between young and old (*P*=0.649), then again being 3% greater in very old compared to young rats (*P*<0.001) (Figure 5D). These more stretched sarcomeres measured in the TA of very old rats, combined with the decreased FL, demonstrated a progressive loss of TA SSN up to very old age: TA SSN decreased 9% from young to old rats (*P*<0.001) and 16% from young to very old rats (*P*<0.001), with notably an 8% decrease from old to very old (*P*=0.048) (Figure 6D).

Collectively, like with the MG, losses of TA muscle mass with age seemed to be driven by losses of both CSA and SSN.

### Changes in vastus lateralis morphology across age groups

Lastly, VL wet weight did not differ between young and middle-aged rats (*P*=0.231) nor between old and very old rats (*P*=0.996), but decreased from young and middle-aged to both old and very old (all *P*<0.001) with an overall decrease of 48% (Figure 3A).

VL ACSA did not differ between young and middle-aged rats (*P*=0.730) nor between old and very old rats (*P*=0.998), but decreased from young and middle-aged to both old and very old rats (all *P*<0.001) with an overall decrease of 49% (Figure 3B). VL PCSA did not differ between young and middle-aged rats (*P*=0.530) nor between old and very old rats (*P*=0.980), but trended toward decreasing from young to old rats (*P*=0.052) and decreased from middle-aged to both old (*P*=0.003) and very old rats (*P*<0.001) with an overall decrease of 35% (Figure 3C).

The VL saw the largest age-related decrease in FL driven by the largest loss of SSN. VL FL did not differ between young and middle-aged rats (*P*=0.999) nor between old and very old rats (*P*=1.00), but decreased from both young and middle-aged rats to old and very old rats (all *P*<0.001) with an overall FL decrease of 19% (Figure 4E). SL also exhibited a 3% increase from young to old rats (*P*=0.018) (Figure 5E) which would further contribute to a lower calculated SSN in old rats. Similar to FL, VL SSN did not differ between young and middle-aged rats (*P*=1.00) nor between old and very old rats (*P*=1.00), but decreased from both young and middle-aged rats to old and very old rats (all *P*<0.001) with an overall SSN loss of 21% (Figure 6E).

Overall, losses of VL muscle mass with age also seemed to be driven by losses of both CSA and SSN.

## Discussion

The Fischer 344/Brown Norway hybrid rat is recommended by the National Institute on Aging as the standard rodent model of skeletal muscle aging due to their ability to reach an advanced age with minimal development of confounding pathologies (21). The present study used this specialized strain to investigate age-related changes in SSN across multiple hindlimb muscles and aging time points, and the contribution to the loss of whole muscle mass. We showed across five hindlimb muscles that the loss of SSN contributes in part to the overall age-related loss of muscle mass in rats, with these SSN losses starting beyond middle age (20 months), though with some muscle-dependent differences. These findings provide key insight into where contractile tissue is lost in muscle during aging, and as SSN is closely tied to biomechanical function (2,22), presents SSN as a distinct therapeutic/rehabilitative target for improving muscle performance in older adults.

### Age-related changes in overall body mass

We observed an initial increase in body mass with age (i.e., from 8 to 20 and 32 months) (Figure 2) despite no changes (from 8 to 20 months) or decreases (from 8 to 32 months) in muscle wet weight (Figure 3). This increase in overall body mass without a corresponding change in muscle wet weight implies a disproportionate increase in non-contractile tissue (e.g., fat mass) over lean mass with age, which is consistent with previous observations (23–26). Body mass also tapered off by 36 months of age such that it did not differ between 8 and 36-month-old rats. This loss of overall body mass at very old age could reflect the greater loss of muscle wet weight from 8 to 36 months than from 8 to 32 months that we observed across most muscles.

### Contribution of serial sarcomere number to the age-related loss of muscle mass

For each of the five muscles included in the present study, no age-related loss of mass (i.e., wet weight) was observed until after 20 months of age (Figure 3A). These results align with previous studies: Rader et al. (27) observed an age-related loss of rat TA mass only at 30 months of age, as compared to 3, 6, and 27 months; Yu et al. (28) observed a loss of rat MG mass starting after 18 months; and Lushaj et al. (24) observed losses of rat VL and vastus medialis muscle mass starting after 21 months. Corresponding to the loss of muscle mass, we also showed here that age-related decreases in SSN, ACSA, and PCSA were not evident until after 20 months of age. This lack of a significant loss of muscle mass from 8 months (a human age of ∼25 years) to 20 months (a human age of ∼50 years) aligns approximately with the trajectory often depicted in humans, with muscle mass plateauing around 20-30 years and experiencing a more accelerated decline starting after 50 years (29,30).

The losses of SSN we observed here in the soleus and MG from 8 to 32 months align with our previous studies on these muscles (4–6), and we can now also expand the age-related loss of SSN to the plantaris, TA, and VL in rats (Figure 6). Furthermore, observing the violin plot shapes in Figure 6, there is notably more variability in SSN across muscle fascicles at 32 and 36 months, suggesting some fascicles lose SSN with aging while others may maintain SSN into advanced age. Sarcomeres contain proteins including (but not limited to) actin, myosin, titin, and α-actinin (which largely comprises the Z-disc) that each have their own mass, therefore the loss of sarcomeres either in the serial or parallel direction across a muscle could amount to loss of mass at the whole-muscle level. To that end, the losses of both SSN and contractile tissue in parallel (represented by PCSA and ACSA) seemed to contribute to the losses of muscle mass across all muscles investigated in the present study. There was, however, some muscle-dependence for how these measures contributed to the age-related loss of muscle mass. For the soleus and tibialis anterior, there were greater losses of SSN and muscle wet weight at 36 months than at 32 months, but only losses of ACSA and PCSA at 36 months. Hence, it appears that age-related losses of mass for the soleus and TA were driven mainly by losses of contractile tissue in series at 32 months and by losses of contractile tissue both in series and in parallel at 36 months. A similar continued loss of soleus wet weight was also observed previously from 31 to 37 months (31). For the plantaris, loss of mass was not evident until 36 months, which corresponded to losses of ACSA and PCSA at 36 but not 32 months. However, a relatively small (7%) loss of SSN in the plantaris was detected at 32 months, which stayed approximately constant at 36 months. Therefore, loss of contractile tissue in parallel seemed to account for more of the loss of mass in the plantaris than contractile tissue in series. Indeed, Blough and Linderman (32) observed loss of plantaris myofibre CSA (a measure of sarcomeres in parallel within a myofibre) for all fibre types alongside decreases in plantaris mass and PCSA in 36-compared to 6-month-old rats. Lastly, for the MG and VL, the losses of contractile tissue in series and in parallel seemed to contribute in tandem to the loss of whole muscle mass.

The age-related loss of contractile tissue in parallel suggested by ACSA and PCSA could reflect two potential scenarios: 1) a decrease in myofibre CSA; or 2) a decrease in the number of myofibres. The present study is limited in that we did not obtain either of these measures, however, a wealth of previous literature provides insight. While studies on humans lean towards the total number of myofibres being a greater contributor than myofibre CSA to the age-related loss of whole muscle ACSA (33–35), studies on rats suggest both could contribute (24,31,32,36–38), with loss of myofibre CSA perhaps contributing more at very old age (i.e., 36 months and beyond) (24). Unlike these more equivocal findings on the loss of myofibre CSA versus myofibre number with age, we show here unequivocally that muscles lose sarcomeres in series with age and that this SSN loss aligns with the loss of a muscle’s total mass.

### Possible driving mechanism for the age-related loss of serial sarcomere number and explanation for muscle-dependent differences

Active lengthening (i.e., eccentric) contractions (5,14) and greater muscle excursions relative to habitual activity (39) stimulate increases in SSN. Along similar lines, when a muscle is forced into a position with shorter SLs relative to its habitual activity, SSN decreases to regulate passive tension and optimize actin-myosin overlap for force production in that new position (40,41). One hallmark of aging in both older humans and rodents is a reduced stride length (42,43). Horner et al. (44) showed in the rat MG that this reduced stride length results in shorter fascicle excursions during locomotion, and speculated that this reduced stride length likely arises from an age-related elevation in passive stiffness in the muscle’s extracellular matrix (45–47). It seems likely, then, that a reduced fascicle excursion with age changes the range of SLs over which the muscle is active, driving the subtraction of sarcomeres in series such as those observed here. Considering these associations between muscle excursion, range of motion, and SSN, it could be speculated that muscles which undergo the longest excursions during locomotion in young healthy rats would exhibit the greatest losses of SSN with aging. Indeed, among the muscles tested, rat VL fascicles have been shown to undergo longer excursions during the stance phase of locomotion (48) than the soleus, MG, and plantaris (49,50), which aligns with the VL undergoing the greatest magnitude (−21%) of SSN loss here. Furthermore, rat plantaris fascicles bear less eccentric strain during locomotion than the soleus and MG (49), which aligns with the plantaris exhibiting the smallest age-related loss of SSN here. MG fascicles also undergo much longer excursions than the soleus during locomotion (49,50), which might seem to go against them having similar magnitude (10-15%) losses of SSN; however, this greater excursion in the MG may explain why it plateaued in its SSN loss by 32 months while the soleus was still undergoing continued SSN loss from 32 to 36 months. Furthermore, since the MG crosses both the knee and ankle, it has a greater possible range of motion than the soleus which only crosses the ankle (51), making the MG potentially face a more drastic change from its habitual activity upon restricting range of motion. As a final note, the age-related loss of SSN does not seem to be fibre type specific, as the muscles investigated here represent a range of fibre type compositions, with the soleus being primarily slow-type, the plantaris, TA, and VL primarily fast-type, and the MG presenting a mixed fibre type (for fibre type compositions see: 52).

### Translatability to humans

Studies comparing old and young healthy humans have reported age-related decreases in FL of 7 to 35% in the VL (53–56), 7 to 19% in the MG (54,57–63), and up to 15% in the soleus (64). The present study shows across five rat hindlimb muscles responsible for ambulation that age-related decreases in FL are driven by losses of SSN, thus, it is possible that these age-related FL decreases reported in studies on humans represent the loss of SSN. However, correct determination of SSN requires measurement of both SL and FL for use in the equation *SSN = FL / SL* like in the present study. As the studies on humans cited above only recorded FL as measured by ultrasound, with no concomitant measurement of SL, it is unclear to what extent those losses of FL represent similar magnitudes of SSN loss. With ultrasound, increases or decreases in the recorded FL could be due to longer or shorter SLs, respectively, at the joint angle in which FL was measured (65). Furthermore, we recently showed that an increase in FL as measured by ultrasound could overestimate the actual increase in SSN (66). Thus, for studies that observed no age-related difference in FL as measured by ultrasound (54,59,60,62,67–73), it is unclear whether this lack of FL difference represents no age-related difference in SSN. Though currently limited in its availability, measurement of SL *in vivo* in humans using microendoscopy has shown that SSN adaptations with training (8,65) and muscle disuse (74) are similar to what has been observed in animals (14,40,41). Thus, we expect age-related decreases in FL in humans are indeed reflective of decreases in SSN like those observed here.

### Methodological considerations

The present study has some limitations to consider. First, due to limitations on available sample size, only male rats were used. The trajectory of the age-related loss of muscle mass and strength in humans is sex-dependent due to the menopausal shift in circulating ovarian hormones in women around age 50 (75), though the age-related hormonal transition in female rats (called “estropause”) is distinctly different from human menopause (76). Collectively, the translatability of our results to female rats is unknown and warrants further investigation. As mentioned, the present study would have also benefitted from measures of myofibre CSA and myofibre number to provide additional insight into changes in contractile tissue arranged in parallel. Our measures of ACSA and PCSA together provided insight at the whole-muscle level and aligned with the abundance of myofibre data from previous studies on aging muscle. With that said, our calculation of PCSA used an assumed muscle density value, and this value may be less applicable to aged muscle which often has greater concentrations of connective tissue and intramuscular fat (77). If greater connective tissue and intramuscular fat were accounted for, the muscle density used for aged muscle could have been lower, leading to greater PCSA values for old and very old rats; thus, our measurements here may represent a conservative (over-)estimate of PCSA’s contribution to age-related muscle mass loss. Lastly, the present study could have benefitted from measures of lean body mass provided by dual-energy X-ray absorptiometry to gain further insight into how our results on the age-related loss of muscle mass from the five muscles investigated align with changes in whole-body lean mass.

## Conclusion

Here we showed that aging results in SSN loss across five rat hindlimb muscles, beginning after middle age and aligning with the loss of whole muscle mass. A muscle’s SSN is closely tied to aspects of its biomechanical performance including the range of motion over which active force can be generated and maximum power production (2,22). Given the unequivocal loss of SSN with aging that we observed here in multiple muscle groups, it is important to develop training interventions to target SSN in aging populations to limit the loss of physical function and maintain independence.

## Acknowledgements

This project was supported by the Natural Sciences and Engineering Research Council of Canada (NSERC). The animals were obtained from the National Institute on Aging (NIA) aged rodent colonies.

## Ethics statement

Approval was given by the University of Guelph’s Animal Care Committee and all protocols followed CCAC guidelines (AUP #4905).

## Data availability

All supporting data are publicly accessible on Figshare at: 10.6084/m9.figshare.28001141

## Author contributions

A.H. and G.A.P. conceived and designed research; A.H. performed experiments; A.H. analyzed data; A.H. and G.A.P. interpreted results of experiments; A.H. prepared figures; A.H. and G.A.P. drafted manuscript; A.H. and G.A.P. edited and revised manuscript; A.H. and G.A.P. approved final version of manuscript.

## References

1. Narici MV, Maffulli N. Sarcopenia: characteristics, mechanisms and functional significance. Br Med Bull. 2010 Sep 1;95(1):139–59.

2. Hinks A, Hawke TJ, Franchi MV, Power GA. The importance of serial sarcomere addition for muscle function and the impact of aging. J Appl Physiol. 2023 Jul 6;135(2):375–93.

3. Hooper AC. Length, diameter and number of ageing skeletal muscle fibres. Gerontology. 1981;27(3):121–6.

4. Power GA, Crooks S, Fletcher JR, Macintosh BR, Herzog W. Age-related reductions in the number of serial sarcomeres contribute to shorter fascicle lengths but not elevated passive tension. J Exp Biol. 2021;224(10):jeb242172.

5. Hinks A, Patterson MA, Njai BS, Power GA. Age-related blunting of serial sarcomerogenesis and mechanical adaptations following 4 weeks of maximal eccentric resistance training. J Appl Physiol. 2024 Mar 21;136(5):1209–25.

6. Hinks A, Power GA. Age-related differences in the loss and recovery of serial sarcomere number following disuse atrophy in rats [Internet]. bioRxiv; 2024 [cited 2024 Jun 12]. p. 2024.06.10.598222. Available from: https://www.biorxiv.org/content/10.1101/2024.06.10.598222v1

7. Lieber RL, Ljung BO, Fridén J. Intraoperative sarcomere length measurements reveal differential design of human wrist extensor muscles. J Exp Biol. 1997 Jan;200(Pt 1):19–25.

8. Andrews MH, S AP, Gurchiek RD, Pincheira PA, Chaudhari AS, Hodges PW, et al. Multiscale hamstring muscle adaptations following 9 weeks of eccentric training. J Sport Health Sci. 2024 Oct 24;100996.

9. Jorgenson KW, Hornberger TA. The Overlooked Role of Fiber Length in Mechanical Load-Induced Growth of Skeletal Muscle: Exerc Sport Sci Rev. 2019 Oct;47(4):258–9.

10. Jorgenson KW, Phillips SM, Hornberger TA. Identifying the Structural Adaptations that Drive the Mechanical Load-Induced Growth of Skeletal Muscle: A Scoping Review. Cells. 2020 Jul 9;9(7):1658.

11. Wroblewski AP, Amati F, Smiley MA, Goodpaster B, Wright V. Chronic exercise preserves lean muscle mass in masters athletes. Phys Sportsmed. 2011 Sep;39(3):172–8.

12. Andreollo NA, Santos EF dos, Araújo MR, Lopes LR. Rat’s age versus human’s age: what is the relationship? Arq Bras Cir Dig ABCD Braz Arch Dig Surg. 2012 Mar;25(1):49–51.

13. Rockenfeller R, Günther M, Clemente CJ, Dick TJM. Rethinking the physiological cross-sectional area of skeletal muscle reveals the mechanical advantage of pennation. R Soc Open Sci. 2024 Sep 18;11(9):240037.

14. Butterfield TA, Leonard TR, Herzog W. Differential serial sarcomere number adaptations in knee extensor muscles of rats is contraction type dependent. J Appl Physiol. 2005;99:7.

15. Hinks A, Jacob K, Mashouri P, Medak KD, Franchi MV, Wright DC, et al. Influence of weighted downhill running training on serial sarcomere number and work loop performance in the rat soleus. Biol Open. 2022 Jul 15;11(7):bio059491.

16. Lieber RL, Yeh Y, Baskin RJ. Sarcomere length determination using laser diffraction. Effect of beam and fiber diameter. Biophys J. 1984 May;45(5):1007–16.

17. Chen J, Mashouri P, Fontyn S, Valvano M, Elliott-Mohamed S, Noonan AM, et al. The influence of training-induced sarcomerogenesis on the history dependence of force. J Exp Biol. 2020 Jun 19;223(Pt 15):jeb218776.

18. Ward SR, Lieber RL. Density and hydration of fresh and fixed human skeletal muscle. J Biomech. 2005 Nov;38(11):2317–20.

19. Zuurbier CJ, Heslinga JW, Lee-de Groot MB, Van der Laarse WJ. Mean sarcomere length-force relationship of rat muscle fibre bundles. J Biomech. 1995 Jan;28(1):83–7.

20. Wilkinson RD, Mazzo MR, Feeney DF. Rethinking the Statistical Analysis of Neuromechanical Data. Exerc Sport Sci Rev. 2023 Jan 1;51(1):43–50.

21. Elliott JE, Omar TS, Mantilla CB, Sieck GC. Diaphragm muscle sarcopenia in Fischer 344 and Brown Norway rats. Exp Physiol. 2016;101(7):883–94.

22. Hinks A, Franchi MV, Power GA. The influence of longitudinal muscle fascicle growth on mechanical function. J Appl Physiol. 2022 Jul 1;133(1):87–103.

23. Feely RS, Larkin LM, Halter JB, Dengel DR. Chemical versus dual energy x-ray absorptiometry for detecting age-associated body compositional changes in male rats. Exp Gerontol. 2000 May;35(3):417.

24. Lushaj EB, Johnson JK, McKenzie D, Aiken JM. Sarcopenia Accelerates at Advanced Ages in Fisher 344×Brown Norway Rats. J Gerontol A Biol Sci Med Sci. 2008 Sep;63(9):921.

25. Mitzelfelt JD, DuPree JP, Seo D oh, Carter CS, Morgan D. Effects of chronic fentanyl administration on physical performance of aged rats. Exp Gerontol. 2010 Oct 15;46(1):65.

26. Gatineau E, Savary-Auzeloux I, Migné C, Polakof S, Dardevet D, Mosoni L. Chronic Intake of Sucrose Accelerates Sarcopenia in Older Male Rats through Alterations in Insulin Sensitivity and Muscle Protein Synthesis. J Nutr. 2015 May;145(5):923–30.

27. Rader EP, Layner K, Triscuit AM, Chetlin RD, Ensey J, Baker BA. Age-dependent Muscle Adaptation after Chronic Stretch-shortening Contractions in Rats. Aging Dis. 2016 Jan;7(1):1–13.

28. Yu BP, Masoro EJ, Murata I, Bertrand HA, Lynd FT. Life span study of SPF Fischer 344 male rats fed ad libitum or restricted diets: longevity, growth, lean body mass and disease. J Gerontol. 1982 Mar;37(2):130–41.

29. McLeod M, Breen L, Hamilton DL, Philp A. Live strong and prosper: the importance of skeletal muscle strength for healthy ageing. Biogerontology. 2016 Jan 20;17:497.

30. Kim KM, Jang HC, Lim S. Differences among skeletal muscle mass indices derived from height-, weight-, and body mass index-adjusted models in assessing sarcopenia. Korean J Intern Med. 2016 Jun 22;31(4):643–50.

31. Fisher JS, Brown M. Immobilization effects on contractile properties of aging rat skeletal muscle. Aging Milan Italy. 1998 Feb;10(1):59–66.

32. Blough ER, Linderman JK. Lack of skeletal muscle hypertrophy in very aged male Fischer 344 × Brown Norway rats. J Appl Physiol. 2000 Apr;88(4):1265–70.

33. Lexell J, Taylor CC, Sjöström M. What is the cause of the ageing atrophy?: Total number, size and proportion of different fiber types studied in whole vastus lateralis muscle from 15- to 83-year-old men. J Neurol Sci. 1988 Apr 1;84(2):275–94.

34. Lexell J. Human aging, muscle mass, and fiber type composition. J Gerontol A Biol Sci Med Sci. 1995 Nov;50 Spec No:11–6.

35. Frontera WR, Reid KF, Phillips EM, Krivickas LS, Hughes VA, Roubenoff R, et al. Muscle fiber size and function in elderly humans: a longitudinal study. J Appl Physiol. 2008 Jun 12;105(2):637.

36. Leeuwenburgh C, Gurley CM, Strotman BA, Dupont-Versteegden EE. Age-related differences in apoptosis with disuse atrophy in soleus muscle. Am J Physiol Regul Integr Comp Physiol. 2005 May;288(5):R1288–1296.

37. Skinner SK, Fenton AI, Konokhova Y, Hepple RT. Variation in muscle and neuromuscular junction morphology between atrophy-resistant and atrophy-prone muscles supports failed re-innervation in aging muscle atrophy. Exp Gerontol. 2021 Dec 1;156:111613.

38. Fuqua JD, Lawrence MM, Hettinger ZR, Borowik AK, Brecheen PL, Szczygiel MM, et al. Impaired proteostatic mechanisms other than decreased protein synthesis limit old skeletal muscle recovery after disuse atrophy. J Cachexia Sarcopenia Muscle. 2023 Jul 14;14(5):2076–89.

39. Koh TJ, Herzog W. Excursion is important in regulating sarcomere number in the growing rabbit tibialis anterior. J Physiol. 1998 Apr 1;508 ( Pt 1)(Pt 1):267–80.

40. Tabary JC, Tabary C, Tardieu C, Tardieu G, Goldspink G. Physiological and structural changes in the cat’s soleus muscle due to immobilization at different lengths by plaster casts. J Physiol. 1972 Jul 1;224(1):231–44.

41. Williams PE, Goldspink G. Changes in sarcomere length and physiological properties in immobilized muscle. J Anat. 1978;127(3):459–68.

42. Kang HG, Dingwell JB. Separating the effects of age and walking speed on gait variability. Gait Posture. 2008 May 1;27(4):572–7.

43. Horner AM, Russ DW, Biknevicius AR. Effects of early-stage aging on locomotor dynamics and hindlimb muscle force production in the rat. J Exp Biol. 2011 Nov 1;214(21):3588–95.

44. Horner AM, Azizi E, Roberts TJ. The interaction of in vivo muscle operating lengths and passive stiffness in rat hindlimbs. J Exp Biol. 2024 Feb 14;jeb.246280.

45. Olson LC, Nguyen TM, Heise RL, Boyan BD, Schwartz Z, McClure MJ. Advanced Glycation End Products Are Retained in Decellularized Muscle Matrix Derived from Aged Skeletal Muscle. Int J Mol Sci. 2021 Aug 17;22(16):8832.

46. Lacraz G, Rouleau AJ, Couture V, Söllrald T, Drouin G, Veillette N, et al. Increased Stiffness in Aged Skeletal Muscle Impairs Muscle Progenitor Cell Proliferative Activity. PloS One. 2015;10(8):e0136217.

47. Kanazawa Y, Miyachi R, Higuchi T, Sato H. Effects of Aging on Collagen in the Skeletal Muscle of Mice. Int J Mol Sci. 2023 Jan;24(17):13121.

48. Gillis GB, Biewener AA. Effects of surface grade on proximal hindlimb muscle strain and activation during rat locomotion. J Appl Physiol. 2002 Nov 1;93(5):1731–43.

49. Hodson-Tole EF, Wakeling JM. The influence of strain and activation on the locomotor function of rat ankle extensor muscles. J Exp Biol. 2010 Jan 15;213(2):318–30.

50. Bernabei M, van Dieën JH, Maas H. Longitudinal and transversal displacements between triceps surae muscles during locomotion of the rat. J Exp Biol. 2017 Feb 15;220(4):537–50.

51. Woittiez RD, Baan GC, Huijing PA, Rozendal RH. Functional characteristics of the calf muscles of the rat. J Morphol. 1985 Jun;184(3):375–87.

52. Armstrong RB, Phelps RO. Muscle fiber type composition of the rat hindlimb. Am J Anat. 1984 Nov;171(3):259–72.

53. Baroni BM, Geremia JM, Rodrigues R, Borges MK, Jinha A, Herzog W, et al. Functional and Morphological Adaptations to Aging in Knee Extensor Muscles of Physically Active Men. J Appl Biomech. 2013 Oct 1;29(5):535–42.

54. Kubo K, Kanehisa H, Azuma K, Ishizu M, Kuno SY, Okada M, et al. Muscle Architectural Characteristics in Young and Elderly Men and Women. Int J Sports Med. 2003 Feb;24(2):125–30.

55. Power GA, Makrakos DP, Rice CL, Vandervoort AA. Enhanced force production in old age is not a far stretch: an investigation of residual force enhancement and muscle architecture. Physiol Rep. 2013 Jun;1(1):e00004.

56. Wu R, Delahunt E, Ditroilo M, Lowery M, De Vito G. Effects of age and sex on neuromuscular-mechanical determinants of muscle strength. Age Dordr Neth. 2016 Jun;38(3):57.

57. Baudry S, Lecoeuvre G, Duchateau J. Age-related changes in the behavior of the muscle-tendon unit of the gastrocnemius medialis during upright stance. J Appl Physiol Bethesda Md 1985. 2012 Jan;112(2):296–304.

58. Conway KA, Franz JR. Shorter gastrocnemius fascicle lengths in older adults associate with worse capacity to enhance push-off intensity in walking. Gait Posture. 2020 Mar;77:89–94.

59. Kelp NY, Gore A, Clemente CJ, Tucker K, Hug F, Dick TJM. Muscle architecture and shape changes in the gastrocnemii of active younger and older adults. J Biomech. 2021 Dec 2;129:110823.

60. Morse CI, Thom JM, Birch KM, Narici MV. Changes in triceps surae muscle architecture with sarcopenia. Acta Physiol Scand. 2005 Mar;183(3):291–8.

61. Narici MV, Maganaris CN, Reeves ND, Capodaglio P. Effect of aging on human muscle architecture. J Appl Physiol. 2003 Dec;95(6):2229–34.

62. Stenroth L, Peltonen J, Cronin NJ, Sipilä S, Finni T. Age-related differences in Achilles tendon properties and triceps surae muscle architecture in vivo. J Appl Physiol Bethesda Md 1985. 2012 Nov;113(10):1537–44.

63. Thom JM, Morse CI., Birch KM, Narici MV. Influence of muscle architecture on the torque and power–velocity characteristics of young and elderly men. Eur J Appl Physiol. 2007 Jun 26;100(5):613–9.

64. Panizzolo FA, Green DJ, Lloyd DG, Maiorana AJ, Rubenson J. Soleus fascicle length changes are conserved between young and old adults at their preferred walking speed. Gait Posture. 2013 Sep;38(4):764–9.

65. Pincheira PA, Boswell MA, Franchi MV, Delp SL, Lichtwark GA. Biceps femoris long head sarcomere and fascicle length adaptations after three weeks of eccentric exercise training. J Sport Health Sci. 2021;11(1):43–9.

66. Hinks A, Franchi MV, Power GA. Ultrasonographic measurements of fascicle length overestimate adaptations in serial sarcomere number. Exp Physiol. 2023 Aug 23;108(10):1308–24.

67. Quinlan JI, Franchi MV, Gharahdaghi N, Badiali F, Francis S, Hale A, et al. Muscle and tendon adaptations to moderate load eccentric vs. concentric resistance exercise in young and older males. GeroScience. 2021;43(4):1567–84.

68. Pinel S, Kelp NY, Bugeja JM, Bolsterlee B, Hug F, Dick TJM. Quantity versus quality: Age-related differences in muscle volume, intramuscular fat, and mechanical properties in the triceps surae. Exp Gerontol. 2021 Dec;156:111594.

69. Barber LA, Barrett RS, Gillett JG, Cresswell AG, Lichtwark GA. Neuromechanical properties of the triceps surae in young and older adults. Exp Gerontol. 2013 Nov;48(11):1147–55.

70. Erskine RM, Tomlinson DJ, Morse CI, Winwood K, Hampson P, Lord JM, et al. The individual and combined effects of obesity- and ageing-induced systemic inflammation on human skeletal muscle properties. Int J Obes 2005. 2017 Jan;41(1):102–11.

71. Franchi MV, Monti E, Carter A, Quinlan JI, Herrod PJJ, Reeves ND, et al. Bouncing Back! Counteracting Muscle Aging With Plyometric Muscle Loading. Front Physiol. 2019 Mar 5;10:178.

72. Gerstner GR, Thompson BJ, Rosenberg JG, Sobolewski EJ, Scharville MJ, Ryan ED. Neural and Muscular Contributions to the Age-Related Reductions in Rapid Strength. Med Sci Sports Exerc. 2017 Jul;49(7):1331–9.

73. Karamanidis K, Arampatzis A. Mechanical and morphological properties of different muscle–tendon units in the lower extremity and running mechanics: effect of aging and physical activity. J Exp Biol. 2005 Oct 15;208(20):3907–23.

74. Adkins AN, Dewald JPA, Garmirian LP, Nelson CM, Murray WM. Serial sarcomere number is substantially decreased within the paretic biceps brachii in individuals with chronic hemiparetic stroke. Proc Natl Acad Sci. 2021 Jun 29;118(26):e2008597118.

75. Samson MM, Meeuwsen IB, Crowe A, Dessens JA, Duursma SA, Verhaar HJ. Relationships between physical performance measures, age, height and body weight in healthy adults. Age Ageing. 2000 May;29(3):235–42.

76. Koebele SV, Bimonte-Nelson HA. Modeling menopause: The utility of rodents in translational behavioral endocrinology research. Maturitas. 2016 Feb 3;87:5.

77. Rivas DA, Morris EP, Fielding RA. Lipogenic regulators are elevated with age and chronic overload in rat skeletal muscle. Acta Physiol Oxf Engl. 2011 Aug;202(4):691–701.

